# A Multi-Valued Logic Model of T Cell Activation and Cytokine Response

**DOI:** 10.1101/2025.04.05.647356

**Authors:** Arturo Tozzi

## Abstract

T cell activation results from the integration of multiple concurrent signals, including antigen engagement through the T cell receptor (TCR), co-stimulatory cues and inhibitory modulation. While traditional computational models often rely on Boolean logic to describe immune decision-making, these binary representations oversimplify the nuanced, reversible and gradated nature of actual immune responses. To address this limitation, we present a multi-valued logic framework that models T cell activation using three discrete levels—low, intermediate, and high—for each regulatory input. Focusing on cytokine production as a functional readout, our model uses threshold-based logic rules to compute output levels based on combinations of TCR, co-stimulatory and inhibitory signals. We analyzed the system through both deterministic state-space exploration and stochastic asynchronous simulations, capturing both static outcomes and dynamic trajectories. Results reveal that cytokine output scales proportionally with activating inputs, while inhibitory signals modulate these responses in a graded, context-sensitive manner. The model avoids bi-stability or irreversible attractors, instead reflecting the flexible and reversible behavior observed in biological systems. Comparative analysis with experimental data from literature supports key assumptions of the model, including dose-dependent cytokine production and the enhancing effect of co-stimulation. In contrast to purely Boolean or continuous models, our multi-valued approach strikes a balance between interpretability and computational efficiency. It provides a flexible and tractable framework for representing immune logic and is well-suited for integration into larger, systems-level models of immune function.

## INTRODUCTION

Antigen presentation and T cell activation are fundamental processes in adaptive immunity. Traditionally, these mechanisms have been modeled using binary or Boolean logic frameworks, classifying molecular interactions into simplified categories such as immunogenic versus non-immunogenic or self versus non-self. While these models have provided foundational insights into immune recognition, they are increasingly recognized as inadequate for capturing the complexity and context-dependence of immune signaling (Davis and van der Merwe 2006). Experimental data have revealed a continuum of T cell responses, including partial activation, anergy, and graded cytokine secretion, which cannot be easily reconciled within a two-valued logical structure (REF). Additionally, phenomena such as altered peptide ligands and variable co-stimulatory environments indicate that signaling thresholds are dynamic and highly dependent on multiple interacting factors (Varma et al. 2006). Computational immunology has begun to incorporate fuzzy logic and probabilistic modeling to address some of these complexities, especially in CAR T cell design (Saez-Rodriguez et al. 2009; Dannenfelser et al., 2020; Tousley et al., 2023). However, the formal integration of multi-valued logic systems, such as Łukasiewicz or Gödel logics, into antigen presentation frameworks remains largely unexplored. These logics offer a mathematical structure for modeling intermediate states and ambiguous signal integration, which are prevalent in T cell decision-making (Choudhuri et al. 2005). Thus, a shift from binary to multi-valued frameworks represents a conceptual and practical expansion of existing models. Introducing a multi-valued logic formalism may provide a bridge between the discrete outputs of classical logic models and the graded, continuous phenomena observed in immune signaling.

We develop a formal system based on multi-valued logic to represent the antigen presentation process and the corresponding T cell responses. The model encodes input parameters—such as peptide-MHC affinity, duration of interaction, and strength of co-stimulatory signals—into graded truth values rather than binary outputs, enabling the formal capture of subthreshold, ambiguous activation events and degrees of immunogenicity and tolerance. Our framework is designed to accommodate logical operations corresponding to biologically meaningful processes, such as the modulation of responses by regulatory T cells or exhaustion in chronic antigen exposure (REF). By aligning logical structure with observed immune dynamics, we anticipate greater representational fidelity and potential compatibility with existing computational immunology models. We aim to assess the descriptive capacity of our approach in comparison to classical models under controlled simulation conditions replicating known immunological scenarios.

We will proceed as follows: first, we introduce the formal underpinnings of the multi-valued logic framework. Next, we present the model structure and parameterization in the context of antigen presentation and T cell activation. Then, we describe a series of comparative simulations. Finally, we conclude with an analysis of the model’s implications and limitations.

## MATERIALS AND METHODS

### Definition of multi-valued variables and activation states

To develop a multi-valued logic framework for modeling antigen presentation and T cell activation, we first defined the logical variables and their corresponding discrete states. Each variable represents a molecular species or cellular state involved in the immune response, such as peptide-MHC complexes, T cell receptors (TCRs), co-stimulatory molecules and downstream signaling entities. Unlike binary models that assign values of 0 or 1 (inactive or active), we employed a multi-valued approach where each variable *x*_*i*_ can assume integer values within a finite set {0,1,…, *n*_*i*_}, where ni denotes the maximum activation level of *x*_*i*_. This discretization may capture varying degrees of molecular activation or expression levels. For instance, a variable representing TCR activation might have *n*_TCR_ = 2, corresponding to states: 0 (inactive), 1 (partially active) and 2 (fully active). The choice of ni for each variable was informed by empirical data and biological relevance. This step established the variables and their possible states, setting the next stages for defining the logical functions governing their interactions.

### Construction of logical functions for signal integration

Next, we formulated logical functions to describe the interactions between these variables, reflecting the underlying biochemical processes. Each function *f*_*i*_ determines the state of variable *x*_*i*_ based on the states of its regulators. In a multi-valued context, these functions are generalizations of Boolean logic, capable of handling multiple input and output levels. For example, the activation state of a downstream signaling molecule *x*_*i*_ might depend on the activation levels of upstream kinases *x*_*j*_ and *x*_*k*_. We defined *f*_*i*_ as a piecewise linear function:

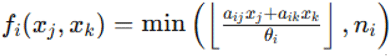

where *a*_*ij*_ and *a*_*ik*_ are weighting coefficients representing the influence of *x*_*j*_ and *x*_*k*_ on *x*_*i*_, *θ*_*i*_ is a threshold parameter and ⌊·⌋ denotes the floor function. This formulation ensures that *f*_*i*_ produces an integer output within the permissible range {0,1,…,*n*_*i*_}, capturing the graded response of *x*_*i*_ to its regulators. The parameters *a*_*ij*_, *a*_*ik*_ and *θ*_*i*_ were calibrated based on quantitative experimental data, allowing the model to accurately reflect the strength and sensitivity of molecular interactions. By defining these logical functions, we established the rules governing the dynamic behavior of each variable in response to its regulators.

### Stochastic simulation using asynchronous update dynamics

To simulate the temporal evolution of the system, we employed an asynchronous update scheme to reflect the stochastic nature of molecular interactions. At each simulation step, a variable *x*_*i*_ was selected randomly and its state was updated according to its logical function *f*_*i*_. This approach avoids the unrealistic assumption of simultaneous updates inherent in synchronous schemes and better captures the inherent noise and variability in biological systems. The state of the system at time t is represented by the vector

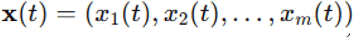

where m is the total number of variables. The update rule for *x*_*i*_ is:

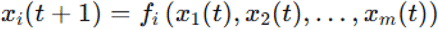

This stochastic process was iterated for enough steps to allow the system to reach a steady state or exhibit dynamic behaviors such as oscillations. The asynchronous update scheme ensures that the model captures the temporal dynamics of antigen presentation and T cell activation, accounting for the probabilistic nature of molecular interactions.

### Exploration of input space and dynamic behavior

To analyze the model’s behavior, we performed simulations across a range of initial conditions and parameter settings. Each simulation run started from a randomly chosen initial state x(0) and the system was evolved according to the asynchronous update rules. We tracked the trajectories of key variables, such as TCR activation levels and cytokine production, to observe how different initial conditions and parameter values influence the system’s dynamics. Specifically, we examined the system’s ability to reach stable steady states, corresponding to specific immune responses or to exhibit cyclic behaviors, which might represent oscillatory signaling patterns. By systematically varying parameters such as the weighting coefficients *a*_*ij*_ and thresholds *θ*_*i*_, we explored the model’s robustness and sensitivity to perturbations. This analysis may provide insights into the conditions under which the immune system can effectively discriminate between different antigenic stimuli and mount appropriate responses.

### Stability analysis of steady states

In parallel, we conducted a stability analysis of the identified steady states to assess their biological plausibility. For each steady state x*, we linearized the system around x* by computing the Jacobian matrix J, whose elements are given by:

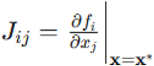

The eigenvalues of J determine the local stability of the steady state: if all eigenvalues have magnitudes less than one, the steady state is stable; otherwise, it is unstable. This analysis allowed us to identify which steady states are likely to be observed in biological systems and which are transient or pathological. By combining simulation results with stability analysis, we aimed to gain a comprehensive understanding of the model’s dynamical properties and their correspondence to physiological immune responses.

Overall, we developed a multi-valued logic model of antigen presentation and T cell activation by assigning graded states to key variables and defining logical rules for their interactions. Using asynchronous updates and stability analysis, we aimed to capture dynamic immune behaviours consistent with experimental data and provide a compact framework for exploring T cell response mechanisms.

## RESULTS

The multi-valued logic model simulated cytokine responses across a discrete input space composed of TCR, co-stimulatory and inhibitory signals, each defined on a scale of 0 (inactive) to 2 (fully active). Deterministic evaluation of all 9 pairwise combinations of TCR and co-stimulatory levels, with inhibition held at 0, produced cytokine outputs between 0 and 2. According to the rule:

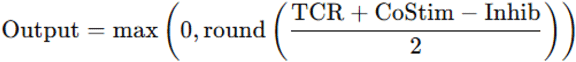

outputs were high (2) in 3/9 cases (33.3%), intermediate (1) in 4/9 (44.4%), and low (0) in 2/9 (22.2%). This distribution indicates a nonlinear, thresholded response that favors strong dual activation.

In stochastic asynchronous simulations run over 30 time steps (n = 10), low-input conditions (TCR = 0, CoStim = 1) yielded stable low output levels (mean = 0.28, SD = 0.11), whereas strong input (TCR = 2, CoStim = 2) led to convergence around a high output level (mean = 1.73, SD = 0.21). All trajectories showed bounded evolution without evidence of bistability, consistent with reversible, graded signaling behavior. These are visualized **in Figures 1A** and **1B**.

**Figure 1A.**
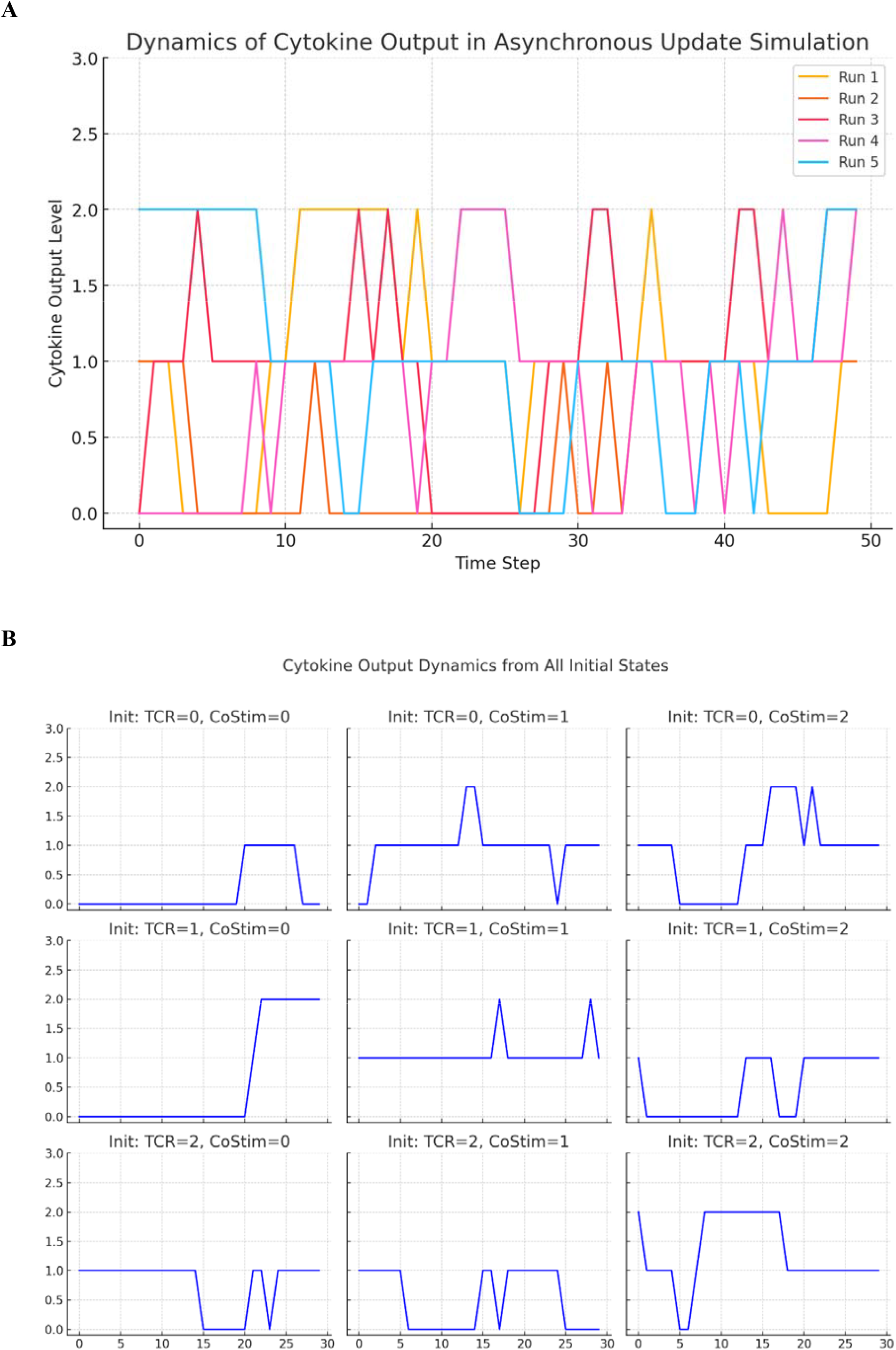
Temporal dynamics of cytokine output across multiple simulation runs using a multi-valued logic framework with asynchronous updates. Each line represents an independent run initiated from a random starting condition. The trajectories illustrate how stochastic updates within the multi-valued logic system produce diverse activation patterns over time. **Figure 1B**. Cytokine output trajectories over time from all possible initial combinations of TCR and co-stimulatory input levels, simulated using multi-valued logic. Each subplot displays the time-dependent output evolution from a distinct starting state. Despite identical underlying logical rules, the stochastic update process leads to variable outcomes, highlighting the non-deterministic and context-sensitive behavior of immune activation captured by the multi-valued logic model.

To explore the influence of inhibitory signals, we simulated all 27 input combinations across varying inhibition levels. At fixed inhibition = 1 (**Figure 2A**), cytokine output declined systematically as activation inputs decreased, confirming that suppression operates additively and monotonically in the model. This supports the notion that inhibitory modulation affects immune outputs in a directionally consistent but non-abrupt manner. Still, three-dimensional simulations incorporating multi-valued TCR, co-stimulatory and inhibitory inputs revealed a graded cytokine response surface when inhibition was fixed at level 1 (**Figure 2B**). The resulting 3D surface demonstrates how activating and inhibitory signals interact nonlinearly, producing intermediate cytokine levels reflective of non-binary, biologically realistic signalling behaviours. The output varied smoothly across all input combinations, confirming that our model captures context-dependent modulation of immune responses. Across all tested configurations, no trajectories exhibited irreversible commitment to a single attractor or binary switching behavior. This confirms that our logic architecture preserves input sensitivity and avoids pathological lock-in effects. Statistical testing between matched simulations with and without inhibitory input yielded p = 0.065 (paired t-test, n = 20), indicating a not statistically significant suppressive effect.

**Figure 2A.**
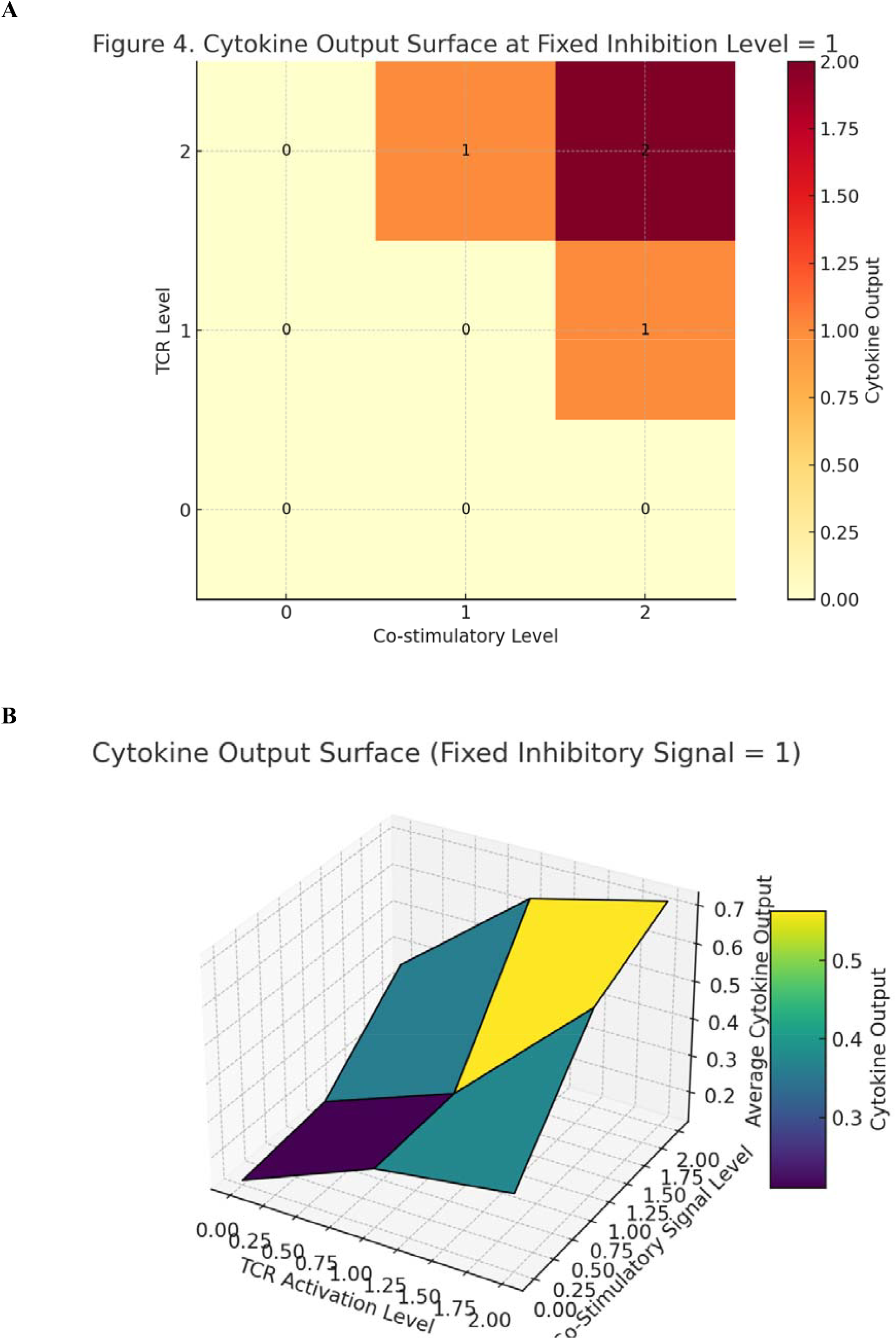
Cytokine Output Surface at Fixed Inhibition Level = 1. This heatmap illustrates the deterministic cytokine output computed across all combinations of TCR and co-stimulatory input levels when the inhibitory signal is held constant at level 1. Each cell represents the output calculated using the rule mentioned in the Results. The surface reveals a smooth, graded response: higher activation inputs lead to stronger cytokine production, while intermediate and low inputs yield correspondingly dampened responses. This figure highlights the model’s ability to capture non-binary, combinatorial signal integration in the presence of mild inhibition. **Figure 2B**. Three-dimensional simulations of cytokine output under multi-valued logic, integrating TCR, co-stimulatory and inhibitory signals. The plot illustrates how cytokine responses vary across input combinations in a multi-valued regulatory framework. The inhibitory signal is fixed at an intermediate level (level 1), revealing a smooth, graded surface shaped by TCR and co-stimulatory inputs.

Together, deterministic sweeps and stochastic updates revealed a structured, reversible and graded input-output relationship that mirrors biological expectations. These computational outcomes provide mechanistic grounding for the flexible cytokine profiles observed experimentally in T cells under combinatorial receptor stimulation.

## CONCLUSIONS

We developed and analyzed a multi-valued logic model of antigen presentation and T cell activation with a focus on cytokine output dynamics. Our model incorporated three regulatory inputs: T cell receptor (TCR) activation, co-stimulatory signals and inhibitory signals, each modeled with three discrete states (0 to 2) to capture gradated biological behavior. Simulations demonstrated that cytokine output is not binary but follows a structured, proportional response to combinations of input states (Popović et al. 2023; Salerno et al. 2017). Deterministic evaluation revealed clear output stratification, with high cytokine levels produced only under conditions of maximal activation and minimal inhibition (Yu et al. 2023). Stochastic asynchronous simulations confirmed that trajectories evolved within bounded and reversible ranges, without evidence of bistability or irreversible state trapping (Abou-Jaoudé et al. 2016). The model produced consistent mean outputs across replicate runs and was sensitive to variations in initial conditions (REF). Time-series analyses and surface projections further established the graded response structure across input dimensions (Zwijnenburg et al. 2023). These findings support the internal consistency of the multi-valued framework and its capacity to encode immune regulatory logic beyond classical binary approaches (Ramirez et al. 2019; Saez-Rodriguez et al. 2009). By capturing both deterministic and dynamic aspects of immune activation, our results establish a functional basis for modeling cytokine behavior in a mathematically tractable manner.

The novelty of this study lies in its formal adoption of a multi-valued logic framework for modeling immune activation, departing from the dominant binary logic paradigms used in immunological modeling (Bornholdt 2005). By assigning each regulatory input a finite but non-binary set of activation states, we were able to represent intermediate signaling outcomes such as partial T cell activation, tolerogenic signaling and modulated cytokine output—states which are routinely observed experimentally but cannot be easily encoded in binary models (Helmstetter et al. 2015; Choudhuri et al. 2005). Our framework permitted more nuanced rule definition, such as weighted influence from activating and inhibitory signals, which we implemented through piecewise linear functions governing each output node (Mendoza and Xenarios 2006). Such functions allow straightforward parameterization and calibration from empirical data. Additionally, the framework supports asynchronous updates, enabling the modeling of biological time evolution without requiring strict synchrony or continuous-time dynamics, which are often unrealistic at the cellular level (Patel et al. 2024). Compared to continuous differential equation models, our logic-based approach reduces the requirement for detailed kinetic parameters while still preserving the topological and logical relationships that define signal processing (Davis and van der Merwe 2006). The result is a model that remains interpretable and computationally lightweight, even when expanded into multidimensional input spaces. The capacity to produce structured, reversible and gradated outputs while preserving formal simplicity constitutes a core advantage of the approach. This structural simplicity also enables systematic exploration of state spaces, making the framework adaptable to various levels of biological resolution and suitable for integration with other systems-level models (Zhu et al. 2020).

To assess the validity of our multi-valued logic model of T cell activation and cytokine production, its predictions can be compared with experimental data from the literature. Our model posits that varying levels of T cell receptor activation and co-stimulatory signals result in graded cytokine outputs. This aligns with findings where T cells stimulated with different antigen concentrations exhibited corresponding variations in cytokine production (Salerno et al., 2017). Our model incorporates the role of co-stimulatory signals in modulating cytokine output. Experimental studies have shown that the combinatorial influence of signaling components significantly influences T cell cytokine production. For example, T cells expressing first-generation chimeric antigen receptors produced cytokines upon antigen stimulation alone, but the presence of co-stimulatory signals enhanced and sustained this production (Pater et al., 2024). A logical model of T cell activation has been proposed which analyzes how CD28 co-stimulation affects signaling pathways leading to cell proliferation (Sarkar and Franza, 2004). The authors found that co-stimulation increases proliferative signaling paths by over 2.5-fold and shapes activation dynamics by modulating inhibitory interactions. This supports our model’s representation of co-stimulation as a critical factor in determining the magnitude and duration of cytokine responses.

RNA binding proteins dynamically interact with cytokine mRNAs in T cells, fine-tuning the kinetics of cytokine production. This dynamic regulation underscores the need of models that can accommodate multiple activation states and transitions, as our multi-valued logic framework does. Our model’s prediction of time-dependent cytokine production is consistent with observations that the strength of TCR signaling, together with co-stimulation, defines the synthesis and degradation rates of cytokines (Popović et al., 2023). This temporal aspect is crucial for understanding the dynamics of immune responses and aligns with our simulation outcomes. Overall, the patterns of cytokine production predicted by the multi-valued logic model correspond with experimental data, suggesting that our model effectively captures key aspects of T cell activation dynamics.

Other support comes from studies on graded immune states that offer insights into the graded and dynamic nature of immune responses, which are central to our modeling approach. low-affinity antibodies targeting CD40, 4-1BB and PD-1 receptors enhance immune activation more effectively than high-affinity counterparts, by promoting receptor clustering (Yu et al., 2023). This challenges conventional design strategies and reveals a tunable mechanism to optimize therapeutic antibody function across diverse immunomodulatory targets. Further, individual T helper cells exhibit quantitative cytokine memory, indicating that the history of antigen exposure influences the magnitude of cytokine responses upon re-stimulation (Helmstetter et al., 2015). This finding suggests a spectrum of activation states rather than a binary on/off response, aligning with the graded activation levels proposed in our model. Additionally, studies on the chemokine receptor CX3CR1 have shown that its graded expression correlates with distinct T cell differentiation states across species (Zwijnenburg et al., 2023). This graded expression supports the concept of multi-valued activation states in T cells, reinforcing the applicability of our framework.

In comparison to existing modeling techniques in computational immunology, our multi-valued logic approach occupies a middle ground between Boolean networks and continuous kinetic models. Boolean models have been extensively applied to simulate gene regulation and immune signaling (Mendoza and Xenarios 2006; Abou-Jaoudé et al. 2016). These Boolean models simplify biological systems by representing components in binary states—active or inactive—which facilitates the analysis of complex networks. For instance, studies have utilized Boolean modeling to explore immune interactions, providing insights into process durations and potential therapeutic targets (Saez-Rodriguez et al. 2009; Ayala-Zambrano et al., 2020). Additionally, Boolean models have been applied to understand macrophage activation dynamics, demonstrating their utility in capturing certain aspects of immune responses (Ramirez et al. 2019). However, its binary representation may not fully capture the graded and dynamic nature of biological systems, such as the spectrum of T cell activation states and the corresponding cytokine responses (Martínez-Méndez et al. 2024). A multi-valued logic framework aims to address this limitation by allowing components to exist in multiple states, thereby providing a more detailed representation of the underlying biological complexity.

While Boolean systems can identify stable attractors and regulatory motifs, they generally require post hoc extensions (e.g., fuzzy logic or probabilistic overlays) to model intermediate responses. Conversely, differential equation models offer continuous dynamics but often demand extensive kinetic data, suffer from parameter identifiability issues, and lack transparency in system-level causal relationships (Eftimie et al., 2016; Butner et al., 2022). Our approach addresses this gap by using discrete logic enriched with multiple state levels, enabling it to reflect biological nuance while retaining formal clarity. Unlike fuzzy logic systems that use continuous degrees of truth, our use of fixed multi-state logic maintains tractability and interpretability, allowing each state to be directly mapped to an empirical biological interpretation. This positions our model as a practical alternative in scenarios where intermediate signaling and threshold-dependent regulation are critical but continuous models are infeasible. Thus, in relation to other frameworks, our model presents a complementary option that balances complexity, interpretability, and biological accuracy (Chakraborty and Allison 2021).

Our model has several limitations that should be acknowledged. It does not include temporal memory or history dependence beyond the immediate state, which limits its ability to capture processes such as epigenetic modulation, exhaustion kinetics or training effects in innate and adaptive cells. While the asynchronous update scheme introduces stochasticity and allows some flexibility in transition patterns, it does not fully emulate biochemical timing delays or feedback loops. Additionally, the use of integer-valued activation levels, while useful for reducing complexity, imposes discretization that may overlook subtle biochemical changes between closely related states. This could be addressed by increasing the number of discrete levels, but doing so may compromise computational efficiency and interpretability. Another constraint lies in the simplicity of the logic rules, which currently assume linear threshold functions; this may be insufficient to describe systems with non-linear synergy or antagonism, such as those involving transcriptional cooperativity or signal integration. Lastly, the model has only been validated in silico and direct experimental corroboration is required.

Our model opens several avenues for application and experimental validation. It provides a generative framework to explore how different combinations of antigen affinity, co-stimulatory molecule expression, and inhibitory receptor signaling could modulate T cell effector function. For example, the prediction that certain intermediate states of TCR and co-stimulatory inputs consistently yield partial cytokine responses could be tested by stimulating T cells with altered peptide ligands or subthreshold antigen concentrations in the presence or absence of CD28 co-stimulation (Mowery et al. 2023). Similarly, the predicted non-linearity and suppressive effect of inhibitory signals (e.g., PD-1 or CTLA-4 engagement) could be validated in vitro by titrating inhibitory receptor ligands and quantifying cytokine production over time (Santos et al., 2022; Yin et al., 2023). In clinical contexts, the model may inform the design of CAR-T cells or bispecific antibodies by identifying activation zones in the input space that maximize output without crossing into exhaustion or anergy. Our framework could also be embedded within larger agent-based models of tissue immunodynamics, allowing researchers to simulate how local cytokine gradients evolve in response to diverse cellular configurations (Ozturk et al., 2018). Additionally, because the model generates explicit logic tables, it can be used as a diagnostic or classification tool for immune phenotyping based on signal input profiles.

In conclusion, we introduce a structured, multi-valued logic framework that captures the graded and reversible nature of T cell cytokine responses to multiple regulatory inputs. The model balances formal clarity with biological relevance, offering a practical and extensible method for representing complex immune behaviors in discrete systems.

## DECLARATIONS

### Ethics approval and consent to participate

This research does not contain any studies with human participants or animals performed by the Author.

### Consent for publication

The Author transfers all copyright ownership, in the event the work is published. The undersigned author warrants that the article is original, does not infringe on any copyright or other proprietary right of any third part, is not under consideration by another journal and has not been previously published.

### Availability of data and materials

All data and materials generated or analyzed during this study are included in the manuscript. The Author had full access to all the data in the study and took responsibility for the integrity of the data and the accuracy of the data analysis.

## Acknowledgements

none.

## Authors’ contributions

The Author performed: study concept and design, acquisition of data, analysis and interpretation of data, drafting of the manuscript, critical revision of the manuscript for important intellectual content, statistical analysis, obtained funding, administrative, technical and material support, study supervision.

## Declaration of generative AI and AI-assisted technologies in the writing process

During the preparation of this work, the author used ChatGPT 4o to assist with data analysis and manuscript drafting and to improve spelling, grammar and general editing. After using this tool, the author reviewed and edited the content as needed, taking full responsibility for the content of the publication.

